# Enamel histomorphometry, growth patterns and developmental trajectories of the first deciduous molar in an Italian early medieval skeletal series

**DOI:** 10.1101/2024.05.13.593149

**Authors:** Stefano Magri, Owen Alexander Higgins, Federico Lugli, Sara Silvestrini, Antonino Vazzana, Luca Bondioli, Alessia Nava, Stefano Benazzi

## Abstract

Understanding the growth patterns and developmental trajectories of teeth during early life stages provides valuable insights into the ontogeny of individuals, particularly in archaeological populations where such information is scarce.

This study focuses on first deciduous molars, specifically investigating crown formation times and daily secretion rates, through histological analysis. A total of 34 teeth from the Early Medieval necropolises of Casalmoro and Guidizzolo (Mantua, Lombardy, northern Italy) were analysed assessing growth parameters and identifying possible differences between sites and between sexes, which are determined through proteomic analysis. Furthermore, a robust linear regression model relating prism length and secretion time was built to estimate growth rates also in teeth in which the finest incremental markings are not visible.

The daily secretion rates (DSR) in inner enamel showed a high homogeneity between dental arches, sexes and the two sites. Values fall within the known range reported in the literature for the same tooth class in archaeological populations. However, a difference in DSR was observed when compared with modern sample published values. Crown formation times and age at crown completion is different between dental arches, with maxillary first molars initiating their matrix apposition earlier than mandibular molars as formerly reported. However, no significant differences were highlighted in association with sex.

This study expands our understanding of the growth and development of first deciduous molars in a medieval population, providing valuable insights into growth trajectories specific to the dental arch. These findings highlight the need for extensive investigations using similar methodologies to attain more accurate and comprehensive information about the developmental patterns of first deciduous molars and underscore the potential use of proteomic analysis to explore deciduous teeth microstructure even in archaeological populations.

## Introduction

Teeth are an invaluable source of information in bioanthropological studies thanks to their remarkable preservation and resistance to diagenetic alterations [1]. They offer insights into various aspects of human biology and culture [2]. The determination of age-at-death [3–6], biological growth patterns [7–10], and dietary habits [11–16] are only a few examples of the valuable information that can be extracted from dental remains. Moreover, the analysis of foetal and infant biological life histories through deciduous dentition offers a profound understanding of populational growth trajectories and health status during early life [9,17,18].

The presence of incremental growth markers in dental enamel, i.e., daily cross-striations and near-weekly – in humans – striae of Retzius [19,20], provides insight into the rhythmical development of this tissue. Specifically, cross-striations are the result of a circadian metabolic variation that occurs during the secretion phase of amelogenesis [21–25] which is associated with differential melatonin production throughout the night-day cycle [26–29]. Cross striations are visible under optical microscopy as an alternation of dark and bright spots along an enamel prism. On the other hand, striae of Retzius, which appear under optical microscopy as dark bands running from the enamel dentine junction (EDJ) to the outer enamel, are the result of slight variations in matrix secretion due to a still debated biorhythm [20,30–36]. Moreover, the identification of stress markers, such as Accentuated Lines (ALs, more pronounced striae of Retzius) and the Neonatal Line (NNL, an AL that forms at birth), allows us to identify respectively the occurrence of physiological stressors during dental development and the birth event [9,37–45]. These latter lines act as permanent records of physiological stress events above a certain threshold – including birth, illnesses, and nutritional deficiencies – which are permanently etched within the enamel’s microstructure during its formation [11,46–49].

By measuring these incremental signatures, we can derive essential parameters such as crown formation times (CFTs; i.e. the time required by ameloblasts - the enamel forming cells - to secrete the whole crown) [50–54], daily secretion rate (DSR; i.e. the daily amount of enamel matrix secreted by an ameloblast) [8,51,55–58], and enamel extension rate (EER; i.e. the speed at which ameloblasts are recruited along the EDJ from apex to neck) [59–61]. These parameters ultimately allow us to reconstruct individual dental crown growth trajectories. In particular, first molars in deciduous dentition offer a glimpse into the growth processes throughout foetal development and the first year of childhood [51,64,65].

Furthermore, the advent of proteomic analysis, particularly the assessment of enamel’s amelogenin, has significantly enhanced our ability to determine the sex of individuals through their dental tissues, even in infant remains and in the case of a single tooth [66–69]. Amelogenin is a key protein coded by a gene whose locus is found on the sex chromosomes [70]. The presence of two different isoforms of this protein, namely AMELX (coded by the gene on the X chromosome) and AMELY (coded by the gene on the Y chromosome), allows for determining the individual’s sex as male. Conversely, the presence of only AMELX enables the estimation of the individual’s sex as female [68]. Until now, osteological sex determination was limited to adult individuals due to the absence of sexual markers on the bones in juveniles. Hence, this analysis expanded our capability to understand sex distribution and variability in populations.

To date, deciduous dentition has not been investigated as profusely as the permanent one [8,18,51,52,58,71–73]. Moreover, among deciduous dentition, many studies have focused on the anterior dental classes [8,10,72]. As for the deciduous molars, most studies have examined variability in modern specimens [54,58], whereas works on archaeological specimens have concerned a few individuals from a specific geographic area [51,65].

The limited number of studies on archaeological deciduous molars, as well as the observation of a slight discrepancy between the development of pre-industrial and post-industrial skeletal remains, highlight the need for further investigations into the variability of – especially – deciduous molars. This can be achieved by expanding the populational coverage at both a diachronic and geographic level to create new population-specific reference standards [5,55,56,74,75].

In this study, we present the histomorphometric analysis of the mesiobuccal cusp of both upper and lower deciduous first molars from inhumed individuals from two Early Medieval Italian necropolises located in the Province of Mantua (Lombardy, northern Italy): Casalmoro and Guidizzolo [76,77]. The Early Middle Ages in the Italian peninsula followed significant social and demographic changes, with the Lombard conquest of Mantua (602 CE) leading to a process of cultural amalgamation, with shifts in socio-economic structures [78]. Within this context, we aim to explore the variability in CFT and enamel growth trajectories by comparing the two sites. Moreover, we expand our known inter-individual variability and investigate the presence of sexual differences in enamel developmental parameters for deciduous first molars. Finally, the data on daily secretion rates was used to develop a new robust regression formula specifically tailored for pre-industrial populations, as previously proposed for deciduous central incisors [8]. This new linear model can be useful to estimate deciduous first molar odontochronologies in cases where daily incremental structures are not visible.

Overall, this study explores the development of prenatal and postnatal enamel of deciduous first molars, advancing our knowledge on the timing and variability of odontogenesis, and providing valuable insights into biological aspects of two Early Medieval populations from northern Italy.

## Materials and Methods

### Archaeological sites and sample

The necropolises of Casalmoro and Guidizzolo, located in the province of Mantua (Lombardy, northern Italy) (Fig 1), were discovered in 1995 and 1996 by the *Soprindentenza Archeologica della Lombardia, Nucleo di Mantova* [76]. The establishment of the two necropolises coincided with the Lombard colonization of northern Italy, which occurred during the 7^th^ and 8^th^ centuries CE [77].

**Fig 1.**
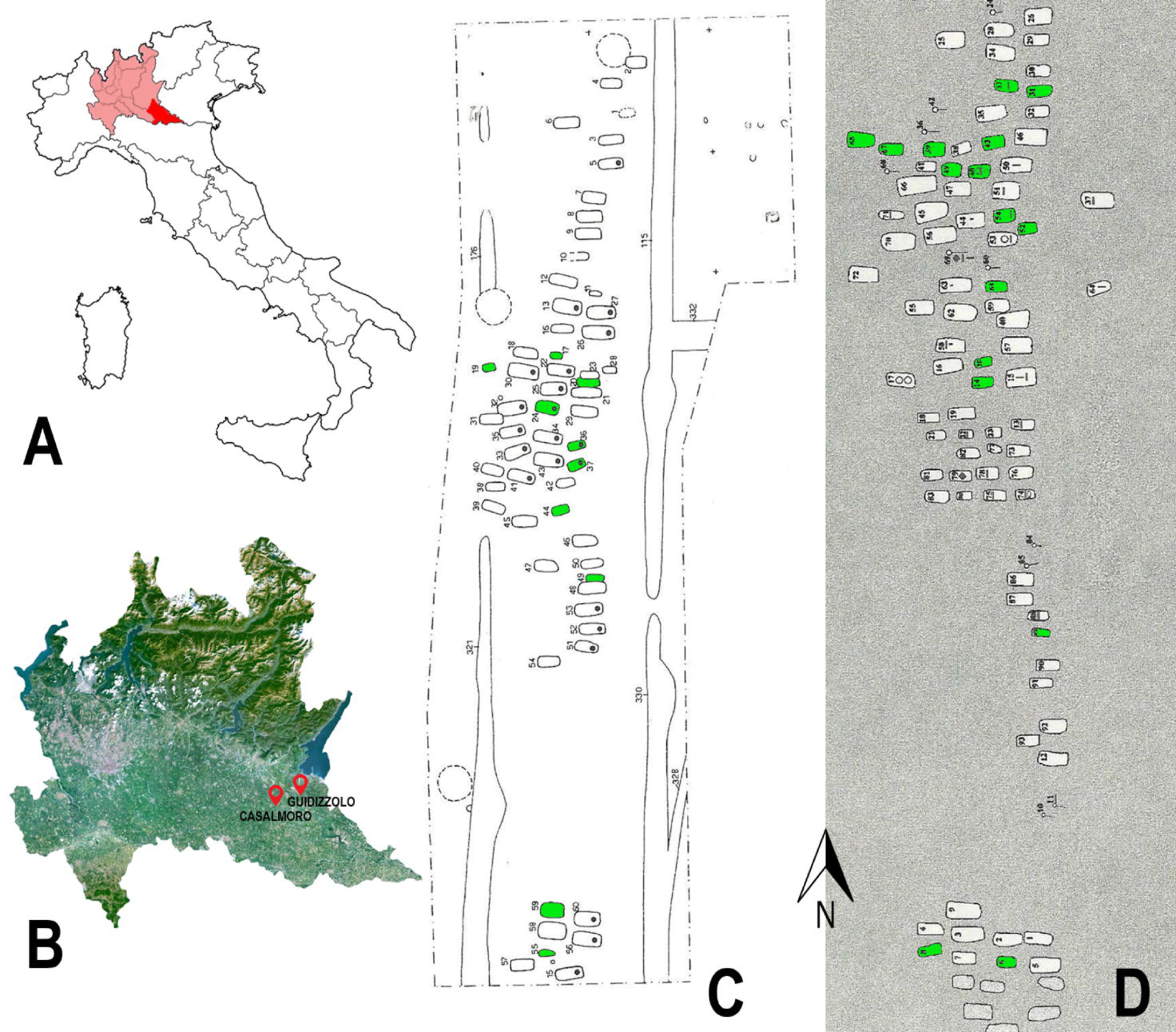
Geographical location and sites plan of the two necropolises. (**A**) Position of the Lombardy region (in light red) and of the province of Mantua (in red) within the Italian peninsula; (**B**) geographic map of Lombardy with the position of the necropolises of Casalmoro and Guidizzolo; (**C**) excavation plan of the necropolis of Casalmoro; (**D**) excavation plan of the necropolis of Guidizzolo (in green: the graves investigated in this study).

Both necropolises were characterized by burials organised in parallel rows, with the human remains – mostly fragmentary and in a poor state of preservation – consistently positioned in a west-to-east orientation (Figs. 1C and 1D), resembling burial practices of contemporary Lombard necropolises [79,80]. The few grave goods found in Casalmoro and Guidizzolo primarily consisted of daggers and bone combs, suggesting a potential association with Germanic populations [76]. Based on the general scarcity of grave goods, Ballarini [76] and Menotti [77] suggested these could be local groups who partially adopted Lombard burial customs. In total, 60 graves were excavated at Casalmoro, whereas 93 were excavated at Guidizzolo. Among these burials, 19 and 25 (from Casalmoro and Guidizzolo, respectively) were infant individuals.

In this study, a total of 34 first deciduous molars (out of 27 individuals) were selected from infants (of unknown sex before proteomic-based sex determination) from both necropolises. Only the deciduous first molars with the crown completely formed were selected. Moreover, the selection was based on cusp and crown preservation and the absence of carious lesions. Specifically, 13 molars (from 10 individuals) were selected from Casalmoro and 21 molars (from 17 individuals) were selected from Guidizzolo. The wear degree for each tooth was evaluated following Molnar [81].

### Sex determination

Enamel samples (approximately 10 mg) were collected from the distolingual cusp of each tooth before histological processing using a precision drill. Each fragment was cleaned with MilliQ (Milli-Q Millipore) water in an ultrasonic bath and then briefly treated with a 5% HCl solution. To extract the peptides, the enamel samples underwent demineralization using a 5% HCl solution for 1 hour. The resulting supernatant was subjected to purification by C_18_ in-house stage tips, followed by elution with 50 μL of 60% acetonitrile in 0.1% formic acid [68,69]. The whole laboratory protocol was performed at the Proteomic facility of the BONES Lab (University of Bologna).

Subsequently, the dehydrated samples were analysed by LC-MS using either the Nano UHPLC Ultimate 3000 coupled to an Exploris™ 480 Hybrid Quadrupole-Orbitrap™ Mass Spectrometer or the Dionex Ultimate 3000 coupled to a HR Q Exactive mass spectrometer (Thermo Scientific), located at Centro Interdipartimentale Grandi Strumenti of the University of Modena and Reggio-Emilia, following Lugli et al. [68] and Granja et al. [82]. Ion chromatograms were manually inspected using Xcalibur^TM^ (Thermo Scientific) and sex estimation followed the presence or absence of peaks associated with the two distinct isoforms of amelogenin that are encoded by the two sexual chromosomes: AMELX-(44-52) ([M + 2H]^2+^ 540.2796 *m/z*) and AMELY-(58-64) ([M + 2H^+^]^2+^ 440.2233 *m/z*) [68,83,84]. No AMEL-peaks were detected in the blank. Raw LC-MS data are available on Zenodo: https://zenodo.org/records/11059821.

### Histological analysis

For histological analysis (following [8]), each tooth was embedded in epoxy resin (EpoThin^TM^ 2, Buehler) and sectioned longitudinally through the mesio-buccal cusps with a microtome (IsoMet^TM^ Low Speed Saw, Buehler) equipped with a 300 μm thick diamond blade. The block containing the dentine horn was gently grinded with P2500 sandpaper (CarbiMet^TM^, Buehler) and polished with 1 μm polycrystalline diamond suspension (MetaDi^TM^ Supreme, Buehler) on a dedicated polishing cloth (TriDent, Buehler) to remove the blade’s scratches. The block was then attached to a glass slide using additional epoxy resin (EpoThin^TM^ 2, Buehler) and sectioned again to obtain a section of ∼300 μm of thickness. Each section was grinded to reach a thickness of ∼150 µm using a sequence of progressive grit sandpapers (CarbiMet^TM^, Buehler) and polished with the 1 μm polycrystalline diamond suspension on the polishing cloth. A thickness of 150 µm was required for future geochemical analyses [7].

Micrographs of the thin sections were taken at 10x magnification using a high-resolution camera (Axiocam 208 color, Zeiss) mounted on a transmitted light microscope (Axioscope 7, Zeiss). The single micrographs were mounted automatically by the tiles feature in Zeiss ZEN core v3.8 software, creating a composite image of each crown section.

Chronologies of crown formation were assessed following the method described in Guatelli-Steinberg et al. [60]. Where the tooth was unworn, the first prism segment measuring 200 μm was drawn from the dentine horn, otherwise from one of the more cuspal prisms available. The stria of Retzius at the end of this first segment was traced down to the EDJ, from which a second 200 μm segment was traced. The procedure was repeated until the neck of the crown (Fig 2). In order to assess the time of the stress events, the chronology of Accentuated Lines was similarly assessed.

**Fig 2.**
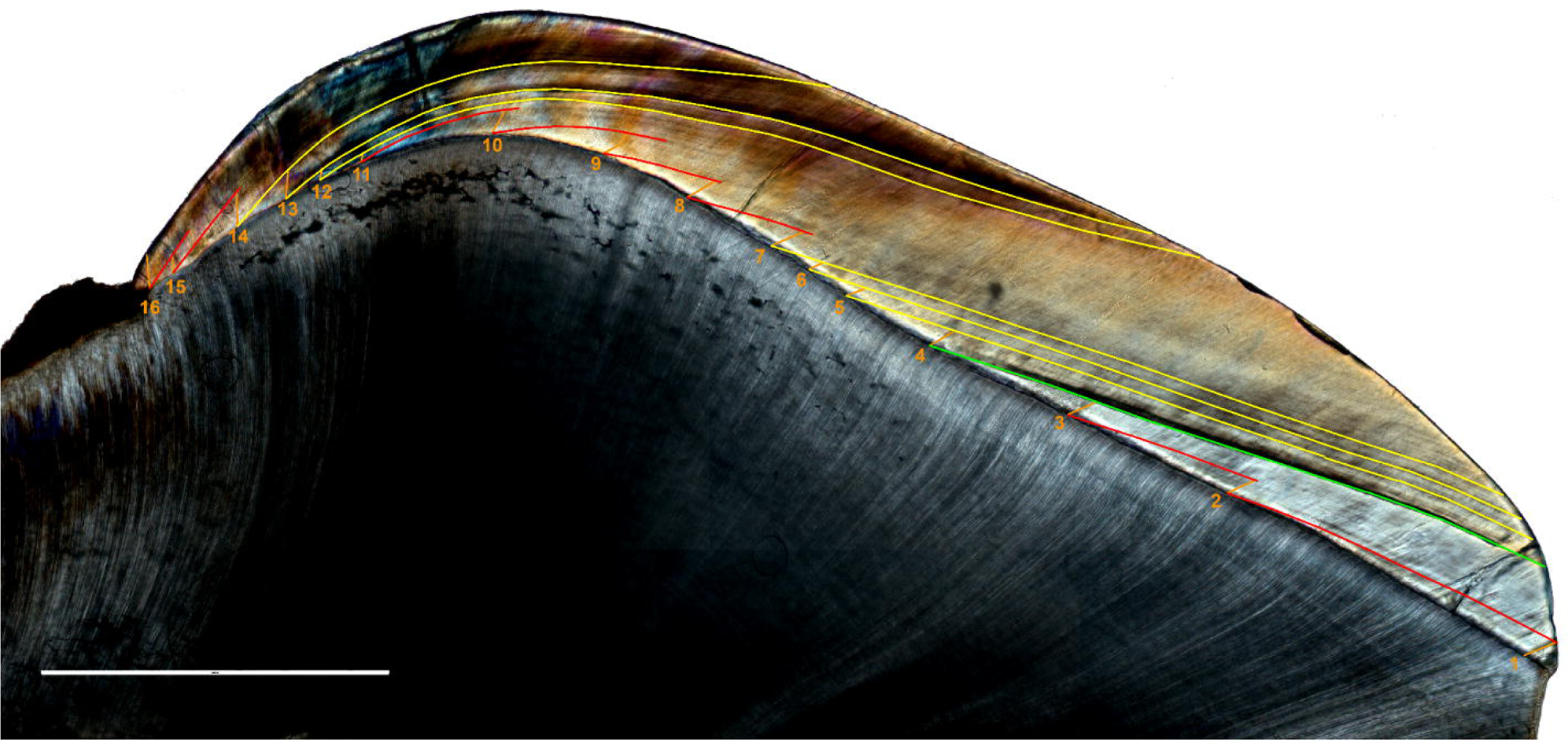
Thin section of Guid T.61 LR buccal side. The Neonatal Line (birth marker) is highlighted in green; Retzius Striae are displayed in red; Accentuated Lines (physiological stressor markers) are showed in yellow; enamel prisms are traced in orange. Scale bar = 1000 µm.

For each prism segment, the length of multiple (n ≥ 6) consecutive cross striations was measured and divided by the corresponding number of cross striations to assess the local mean daily secretion rate (DSR). To estimate the number of days represented by each prism segment, its length was divided by the local mean DSR. The number of days represented by each prism segment was cumulatively summed to obtain the overall crown formation time (CFT). Moreover, by using the Neonatal Line (NNL) as a chronological marker, we distinguished between prenatal and postnatal crown formation, identifying crown initiation (Ci) and crown completion (Cc) times. To estimate average CFTs, teeth lacking the cervical portion (n = 2) were excluded, as they provide partial information. Similarly, also teeth which present a worn dentine horn (n = 22) were omitted in the CFT and Ci estimation.

Local enamel extension rates were assessed by dividing the length of the EDJ by the estimated number of days.

All lines were traced, and measured with ImageJ (v1.54, National Institute of Health, USA) software [85].

Distances were plotted against cross-striation counts to derive the regression formula for the deciduous first molars. In order to compensate for the presence of possible outliers, a robust regression method was adopted [8], with the constraint of the intercept equal to zero (no prism length equal to no days of enamel matrix production) [86–88].

To observe the relationship between EER and days of formation, a Generalized Additive Model (GAM) [87] was adopted. The model was created as a thin-plate spline interpolation on the dependent variable with a number of degrees of freedom which minimizes the error and the residual deviance.

Five teeth were selected to observe the DSRs (n = ∼ 400) throughout the entire buccal crown area and were intentionally selected to provide comprehensive representation across both dental arches and sexes following Nava et al. [8] and Peripoli et al. [10]. The topographical maps of the variation in enamel secretion rates were obtained employing a GAM surface interpolation [87]. Indeed, the maps were created by interpolating DSR values across the enamel area, defined by the local coordinates of the sampling points, and digitized through the software tpsDig v.2.05 software [89].

All statistical analyses were carried out using the program for statistical computing R (v.4.3.2) [90] with mgcv [91], robustbase [88], and lava [92] packages. The distribution of values for the various crown parameters was assessed using a Shapiro-Wilk normality test. If the distribution was found to be normal (p-value > 0.05), group variances were compared using an F-test before applying a parametrical test to evaluate the differences. Conversely, if the values did not follow a normal distribution (p-value < 0.05), a non-parametric test was employed to assess the differences.

## Results

Proteomic analysis estimated a total of 8 females and 19 males within the sample. Among the individuals from Casalmoro, 3 were identified as female and 7 as male (S1 Fig), whereas in the sample from Guidizzolo, 5 females and 12 males were identified (S2 Fig). A comprehensive overview of the enamel developmental parameters, wear stage of the mesio-buccal cusp, and estimated sex for each individual is presented in Table 1.

**Table 1.** Crown formation times parameters in days, sex, number of accentuated lines and sex of each specimen. CFT = Crown Formation Time; CI = Crown Initiation; CC = Crown completion; DSR = Daily Secretion Rate; LL = lower left; LR = lower right; UL = upper left; UR = upper right. ^a^ Teeth used for CFT and CI. ^b^ Teeth excluded for CC. ^c^ Values are calculated based on teeth with an unworn dentine horn or/and an unbroken cervical portion of the crown.

| ID | Dental Arch | Sex | Wear stage<br>(*Molnar, 1971*) | CFT (days) | CI (days before birth) | CC (days) | DSR (µm/day) | Accentuated Lines (n) |
| --- | --- | --- | --- | --- | --- | --- | --- | --- |
| <sup>a</sup>CA T.17 LL | Lower | M | 1 | 418 | 199 | 219 | 3.05 | 3 |
| CA T.17 UR | Upper |  | 1 | 367 | 174 | 193 | 3.18 | 1 |
| <sup>a</sup>CA T.19 UL | Upper | M | 1 | 353 | 171 | 182 | 3.16 | 1 |
| CA T.19 LL | Lower |  | 1 | 401 | 175 | 226 | 3.25 | 2 |
| CA T.20 UL | Upper | M | 3 | 386 | 176 | 210 | 3.18 | 1 |
| CA T.24 LR | Lower | M | 4 | 404 | 149 | 255 | 3.10 | 2 |
| <sup>a</sup>CA T.36 UL | Upper | M | 1 | 358 | 165 | 193 | 3.23 | 1 |
| CA T.37 UR | Upper | M | 2 | 369 | 198 | 171 | 3.24 | 0 |
| CA T.44 UL | Upper | M | 3 | 374 | 180 | 194 | 3.23 | 0 |
| <sup>a</sup>CA T.55 LL | Lower | F | 1 | 401 | 175 | 226 | 3.10 | 0 |
| CA T.59a LR | Lower | F | 2 | 390 | 182 | 208 | 3.19 | 1 |
| CA T.59a UL | Upper |  | 3 | 369 | 181 | 188 | 3.27 | 0 |
| <sup>a</sup>CA T.59b UL | Upper | F | 3 | 343 | 166 | 177 | 3.17 | 0 |
| GUID T.6 LR | Lower | F | 1 | 418 | 185 | 233 | 3.15 | 0 |
| <sup>a</sup>GUID T.8 LR | Lower | M | 4 | 392 | 135 | 257 | 3.12 | 1 |
| <sup>a</sup>GUID T.14 LR | Lower | M | 2 | 418 | 175 | 243 | 3.08 | 0 |
| GUID T.20 UR | Upper | M | 2 | 356 | 181 | 175 | 3.21 | 0 |
| <sup>a</sup>GUID T.31 LR | Lower | F | 3 | 389 | 165 | 224 | 3.11 | 4 |
| <sup>b</sup>GUID T.33 UR | Upper | F | 3 | 307 | 195 | 112 | 3.19 | 0 |
| <sup>a</sup>GUID T.39 LR | Lower | M | 1 | 410 | 179 | 231 | 3.24 | 1 |
| <sup>a</sup>GUID T.43 LR | Lower | M | 2 | 383 | 150 | 233 | 3.21 | 0 |
| GUID T.43 UR | Upper |  | 3 | 359 | 160 | 199 | 3.18 | 0 |
| GUID T.48 LR | Lower | F | 2 | 388 | 169 | 219 | 3.24 | 0 |
| <sup>a</sup>GUID T.49 LR | Lower | M | 2 | 411 | 187 | 224 | 3.18 | 3 |
| <sup>a</sup>GUID T.49 UL | Upper |  | 3 | 357 | 201 | 156 | 3.11 | 0 |
| GUID T.49b LL | Lower | M | 2 | 408 | 178 | 230 | 3.14 | 3 |
| GUID T.52 LL | Lower | M | 2 | 378 | 185 | 193 | 3.13 | 1 |
| <sup>b</sup>GUID T.54 UL | Upper | M | 2 | n.a. | 185 | n.a. | 3.23 | 0 |
| GUID T.61 LR | Lower | F | 5 | 398 | 88 | 310 | 3.07 | 6 |
| GUID T.65 LR | Lower | M | 2 | 383 | 172 | 211 | 3.17 | 1 |
| GUID T.65 UR | Upper |  | 3 | 344 | 187 | 157 | 3.15 | 2 |
| GUID T.67 LL | Lower | M | 3 | 383 | 164 | 219 | 3.19 | 0 |
| GUID T.67 UL | Upper |  | 3 | 325 | 172 | 153 | 3.18 | 0 |
| GUID T.89 UL | Upper | M | 3 | 350 | 195 | 155 | 3.14 | 4 |
| MEAN |  |  |  | 386 (sd = 27)<sup>c</sup> | 172 (sd = 19)<sup>c</sup> | 208 (sd = 34)<sup>c</sup> | 3.17 (sd = 0.26) |  |

The overall mean inner enamel (within 200 µm) daily secretion rate – estimated at 3.17 μmday^−1^ (n = 1855, sd = 0.26) – and the minimum and maximum values recorded are reported in Table 2. The prenatal portions of the crowns’ buccal aspect exhibit a mean inner enamel DSR of 3.20 μmday^−1^, whereas the postnatal portions of the same aspect have a mean inner enamel DSR estimated at 3.12 μmday^−1^ (Table 2).

**Table 2.** Daily secretion rate values in the inner enamel. Minimum, maximum, and mean DSRs value (µm/day) within 200 μm from the EDJ.

|  | n | Mean | sd | Min | Max |
| --- | --- | --- | --- | --- | --- |
| total | 1855 | 3.17 | 0.26 | 2.32 | 4.33 |
| prenatal | 1002 | 3.20 | 0.26 | 2.48 | 4.33 |
| postnatal | 853 | 3.12 | 0.25 | 2.32 | 4.03 |

Among Casalmoro’s specimens, the mean DSR within inner enamel was estimated at 3.18 μmday^−1^ (sd = 0.27, n = 706), whereas it was estimated at 3.16 μmday^−1^ (sd = 0.25, n = 1093) for Guidizzolo (Mann-Whitney U test W = 403736, p-value > 0.05). A significant difference arises when comparing prenatal and postnatal portions across both sites (Mann-Whtiney U test W = 502743, p-value < 0.05). However, the relatively small effect size (Cohen’s d test, d = 0.32) suggests that the observed difference may not be practically meaningful. The similarities between the two sites are also evident in the topographic maps of DSR variation across the buccal aspects of five selected specimens (Fig 3). The maps, as expected, exhibited the lowest values in the inner enamel, with DSR values increasing towards the enamel surface, showing an overall similar pattern in all samples.

**Fig 3.**
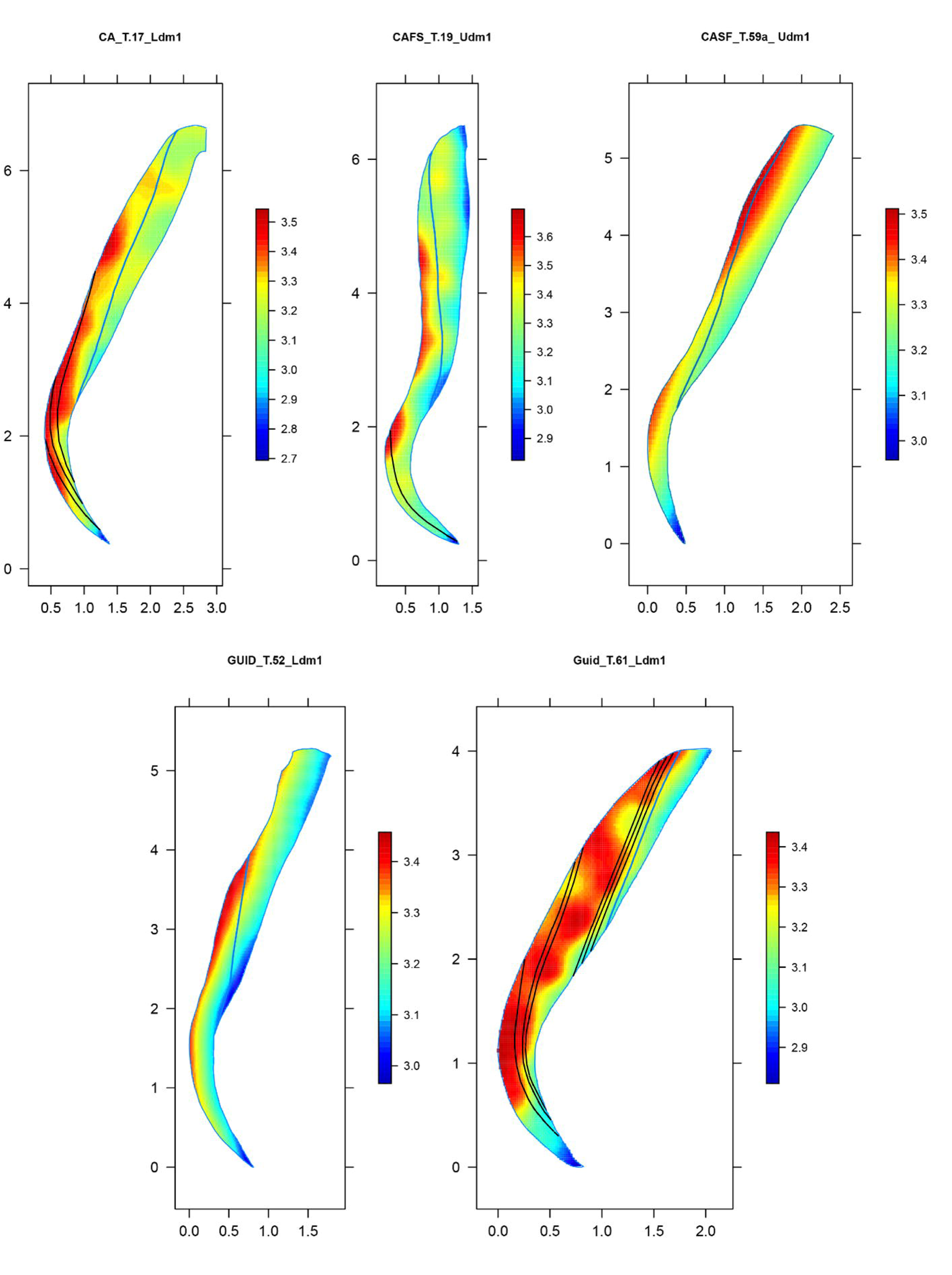
Maps of the topographic distribution of daily secretion rates in the whole mesio-buccal enamel of five individuals. X and Y axes report the topographic coordinates; colour bar = DSR value.

Overall mean CFT was estimated at 386 days (sd = 27, n = 12). A difference of ca. 50 days was observed when comparing dental arches (independent t-test for two samples with equal variances F = 0.26; t = −6.75, df = 10, p-value < 0.05). The mean CFT for upper first molars was estimated at 353 days (sd = 7, n = 5), whereas mean CFT for lower first molars was estimated at 403 days (sd = 14, n = 7). No significant differences were observed between sexes (independent t-test for two samples with equal variances F = 1.25; t = 0.61, df = 10, p-value > 0.05) (Fig 4), nor between the sites (t-test for two independent samples with equal variances F = 0.41; t =0.40, df = 10, p-value > 0.05).

**Fig 4.**
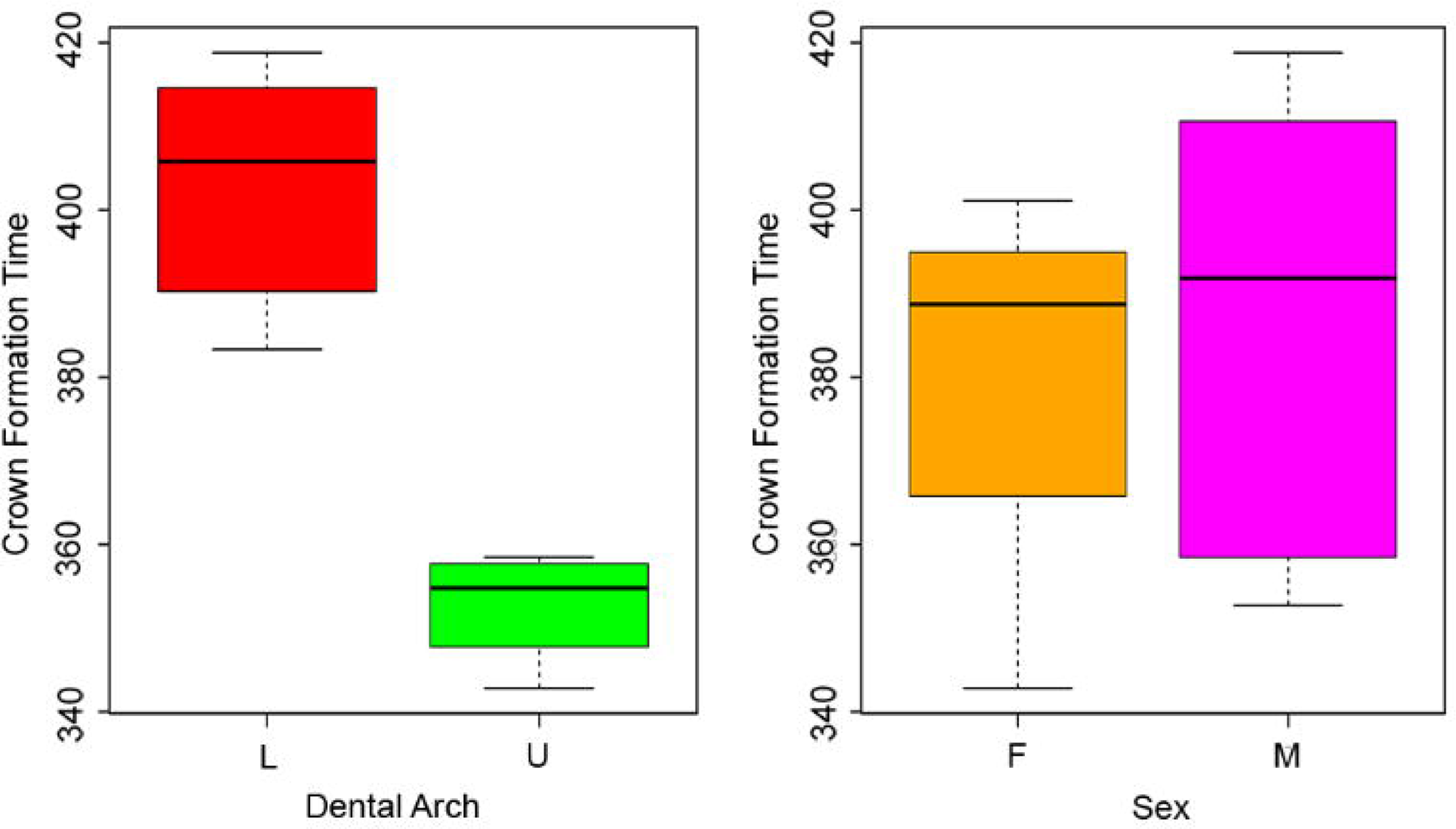
Boxplots of the variation of Crown Formation Time. Boxplots show the variation of CFT depending on arches and sexes.

Mean Ci across the whole sample was estimated at 172 days before birth (sd = 19, n = 12). Considering the different dental arches, mean Ci was estimated at 176 days (sd = 17, n = 5) for upper deciduous molars, and 171 days (sd = 20, n = 7) for lower deciduous molars. This difference is not statistically significant (Mann-Whitnhey U test W = 17, p-value > 0.05), and no significant dissimilarities are observed between males and females (independent t-test for two samples with equal variances F = 14.29, t = 0.78, df = 4.96, p-value > 0.05) (Fig 5).

**Fig 5.**
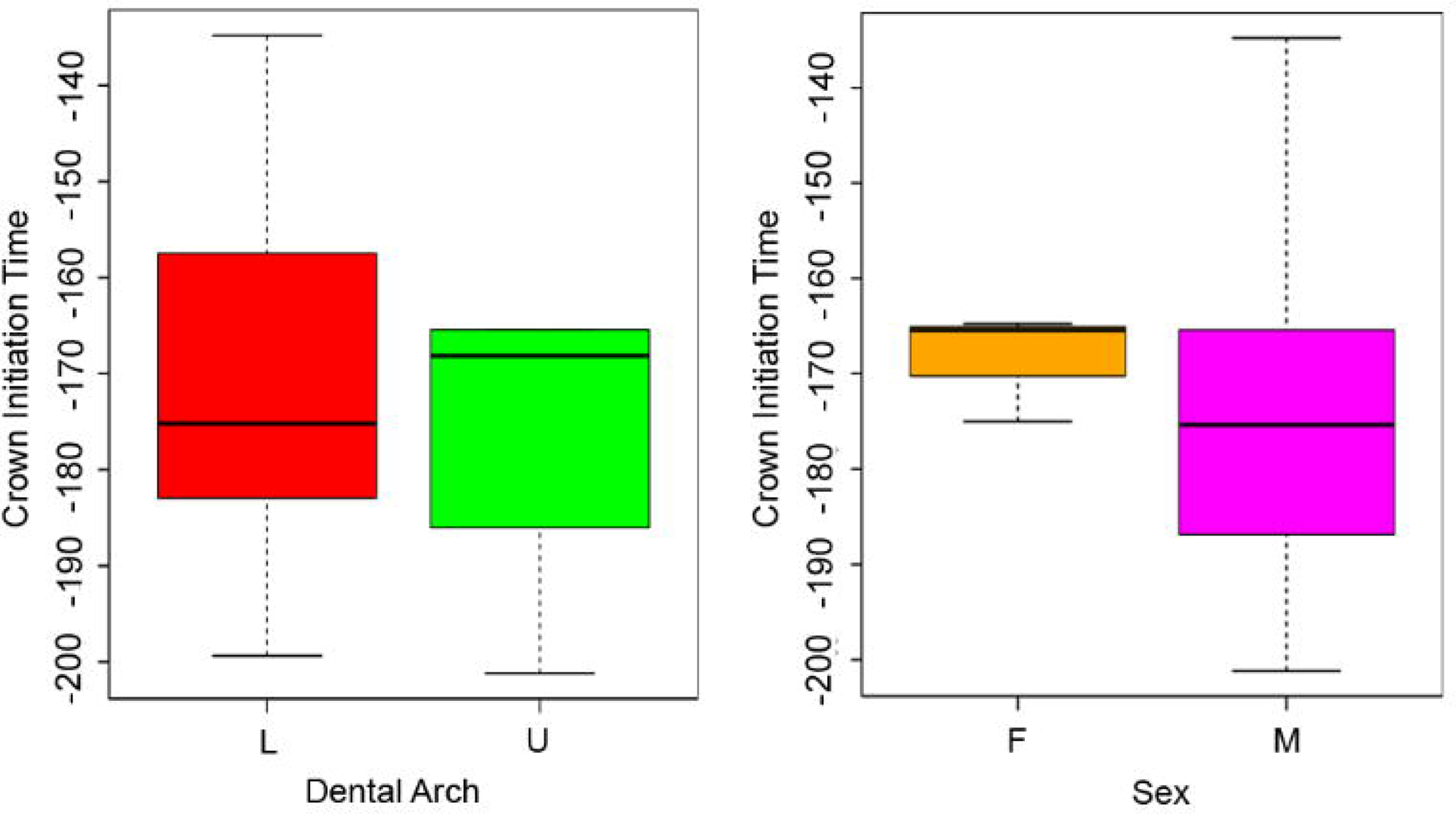
Boxplots of the variation of Crown initiation. Boxplots show the variation of Ci depending on arches and sexes.

Mean age at Cc across the whole sample was estimated at 208 days (sd = 34, n = 32) after birth. Comparing dental arches, lower first molar crowns are completed later than upper molars, with a mean Cc of 231 days (sd = 25, n = 18) after birth, compared to 178 days (sd = 18, n = 14) after birth for the upper molars (Mann-Whitney U test W = 247, p-value < 0.05) (Fig 6). When sex is taken into consideration, mean Cc is estimated at 203 days (sd = 32, n = 24) and 223 days (sd = 40, n = 8) after birth in males and females, respectively, with no statistically significant difference (t-test for two independent samples with equal variances F = 0.62, t = −1.43, df = 30, p-value > 0.05) (Fig 6).

**Fig 6.**
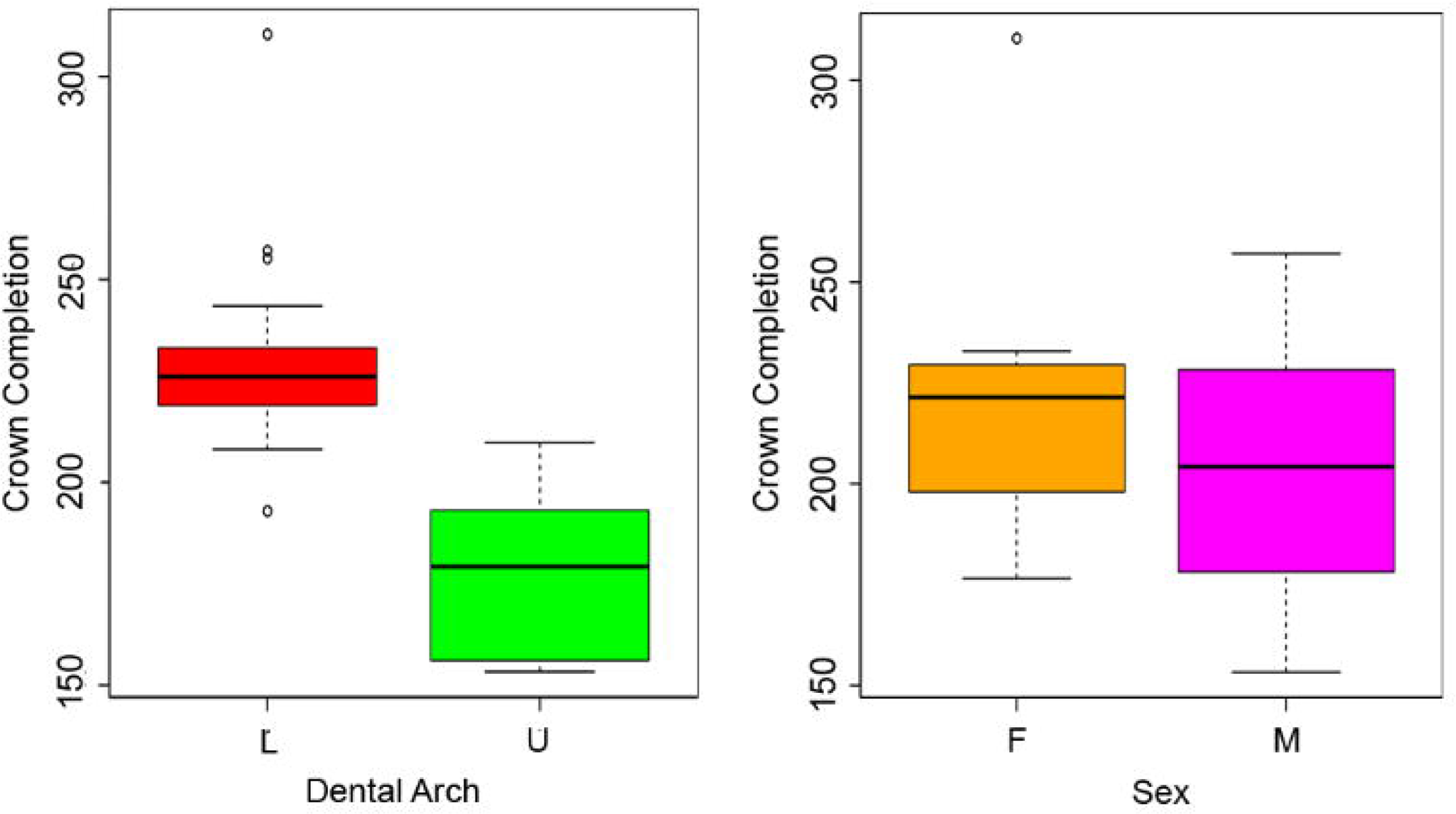
Boxplots of the variation of Crown completion. Boxplots show the variation of Cc depending on arches and sexes.

Enamel extension rate (EER) trends along the crowns’ longitudinal growth are illustrated in Fig 7. Irrespective of the dental arch, the trends along the enamel-dentine junction (EDJ) remain relatively consistent. As expected, ameloblasts’ recruitment is rapid in the cuspal portion and progressively decreases in speed until birth. In postnatal enamel, the speed remains relatively constant, with slight changes coinciding with the presence of Accentuated Lines (ALs) (S4 Fig). Finally, as expected, the speed of ameloblast recruitment decreases to its lowest in proximity of the cervical area.

**Fig 7.**
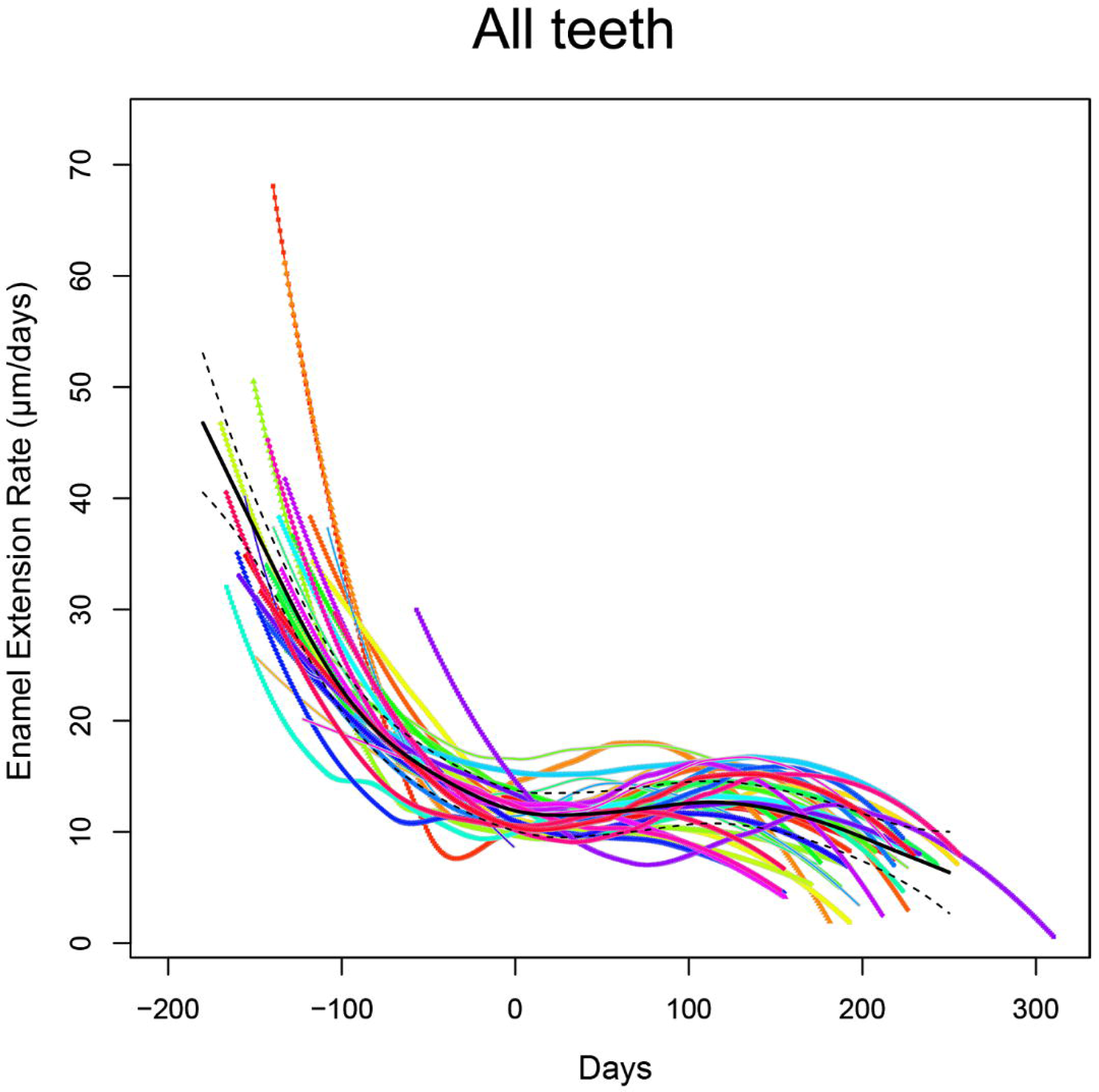
Enamel Extension Rate trend across the crown chronologies. EER trend is represented by a generalized additive model, built as a thin-plate spline interpolation. The black line shows the mean trend of EER in the entire sample.

The robust linear regression model (Fig 8) formulated specifically for first deciduous molars, following Nava et al. [8] work on deciduous incisors, can be represented as Y = 0.316 X (Y = prism lengths; X = days) with an adjusted R^2^ equal to 0.998 to be considered as indicative only because the intercept was forced to be zero [8].

**Fig 8.**
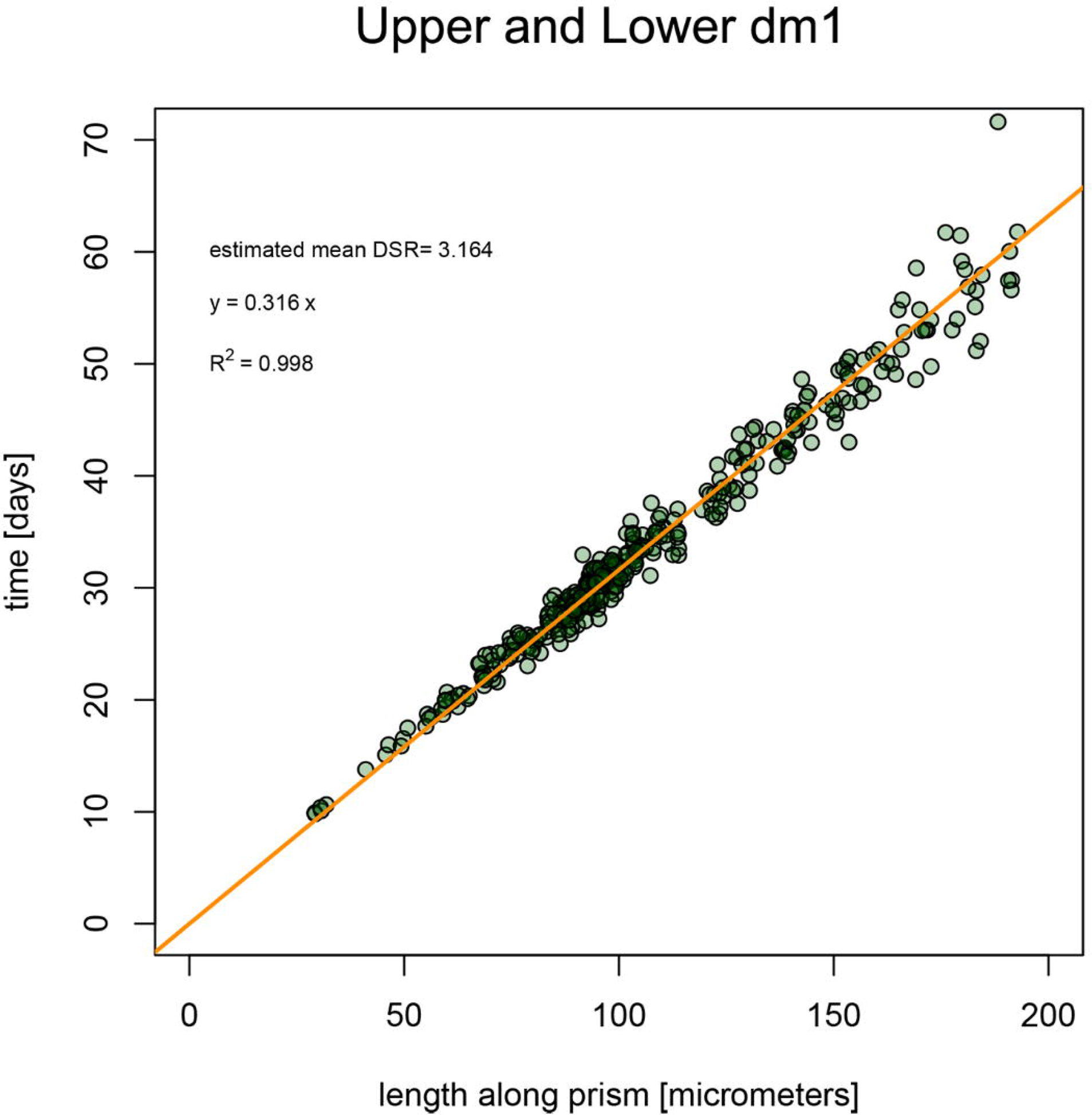
Linear model to estimate the crown formation time from prism lengths on first deciduous molars. The regression formula is Y = 0.316 X (Y = prismal lengths; Y = days) with an adjusted R^2^ equal to 0.998.

## Discussion

Acquiring detailed information on the timing of dental formation is crucial for estimating growth rates and ontogenetic trajectories during the early stages of life, as proxied by dental growth. This is particularly critical when dealing with archaeological populations, for which such data are more challenging to obtain.

In this study, which entails a comparative analysis of two necropolises from the same period (Early Middle Ages) and geographic area (ca 15 km apart), we observe a significant homogeneity in inner enamel DSRs (Fig 8 and Fig 9).

**Fig 9.**
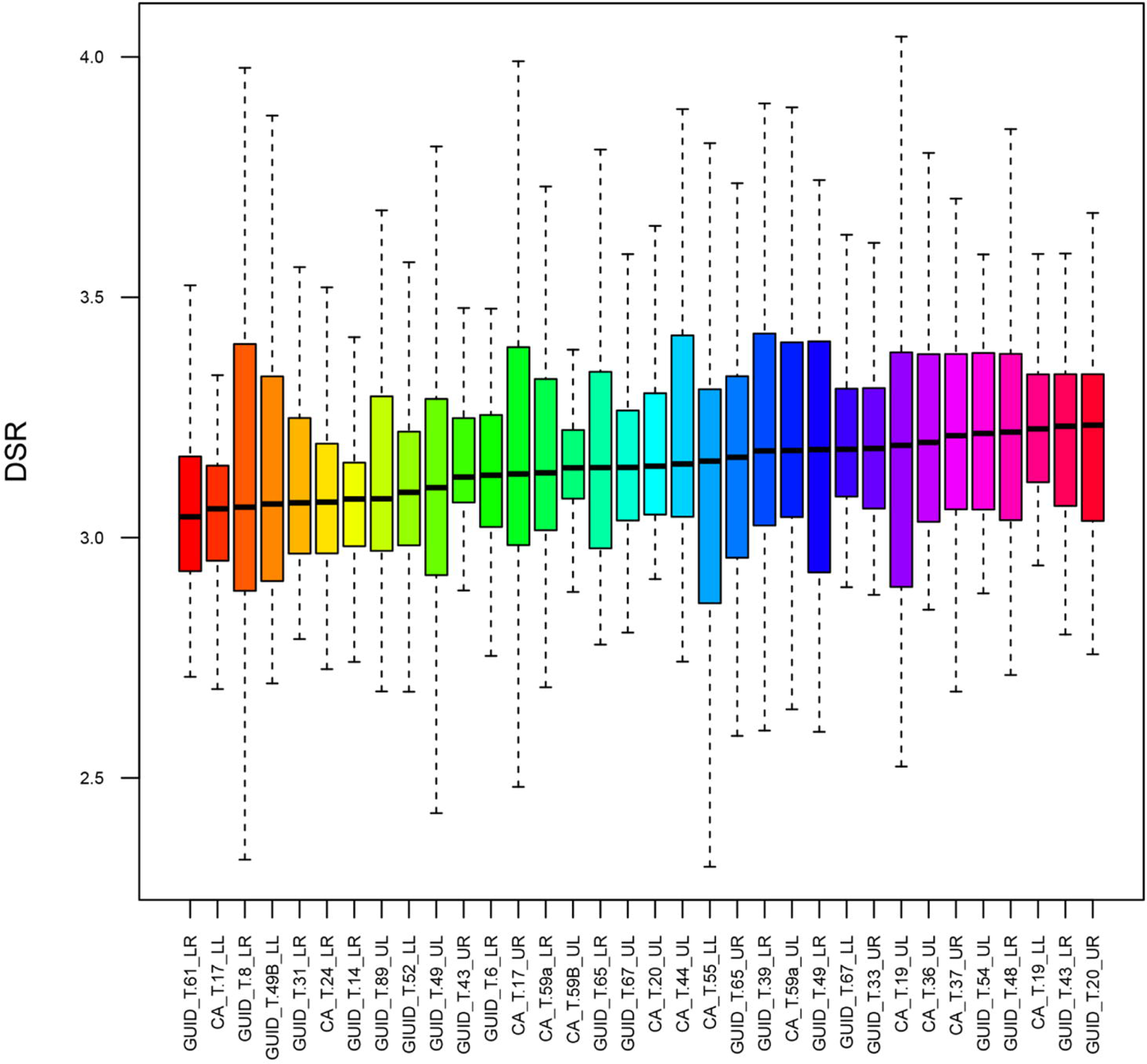
Boxplots of inner enamel DSR values for each tooth. Boxplots show the DSR distribution in the entire sample, ordered by increasing median values.

The values observed in this study indicate a lower inner enamel daily secretion rate in the deciduous molars of this medieval cohort compared to reported modern populations [58]. This finding is in opposition to the results of Aris [56] regarding permanent molars, as well as those of Aris et al. [55] and Nava et al. [8] for deciduous anterior teeth. These works reported a decrease in enamel DSRs over time, suggesting a trend towards a slower developmental pace in human recent evolution. Anyhow, it is worth noting that McFarlane et al. [58] did not differentiate between first and second deciduous molars, which could have potentially influenced the DSR modern comparative values. Overall, the trend of daily secretion rates throughout the entire buccal aspect is consistent with previous observations on deciduous molars [51,65,71]. The lowest values are consistently observed in the inner enamel along the EDJ, with a slight further decrease around the cuspal region, the Neonatal Line and the cervical portion (Fig 3). When comparing the daily secretion maps with those presented in earlier studies [8,10] the distribution trend remains similar to that recorded in anterior teeth, albeit with generally lower DSRs compared to incisors and canines. However, an exception is noted in the case of Guid T.61 LR, which exhibits a more irregular pattern, potentially linked to the presence of a significant number of ALs (n = 6) in postnatal enamel, as ALs represent stressful events in an individual’s life that affect ameloblast activity [21] (Fig 2).

The linear regression models of the upper and lower arches (Fig 8. S3) confirms a consistency in DSRs within inner enamel and appears similar to previously regressed models [8,54]. The slope of this new model is comparable to the one published by Birch and Dean [54], but the intercept is forced to 0, reflecting that no enamel is secreted at day zero.

The mean time of crown formation for the lower first deciduous molars, estimated at 403 days, aligns closely with the findings of Mahoney [51], who reported an average CFT of 388 days on a sample of 12 first deciduous molars. However, our result differs notably from the average CFT of 326 reported by Birch and Dean [54]. This discrepancy with the results of Birch and Dean [54] may however be attributed to the broad variability observed in human populations. In contrast, the estimated mean CFT for upper molars, calculated at 353 days, shows a deviation from the findings of Mahoney [65], who reported a mean CFT of 398 days. Nonetheless, our results are within the range of variability noted in the latter work, which reported CFTs ranging from 336 to 510 days for upper deciduous first molars. The higher CFT identified in lower molars compared to upper molars (as illustrated in Fig 3) can be attributed to the greater length of the EDJ, as EERs are comparable (Fig 7). Additionally, the divergence in CFTs observed between the two sites (Fig 10B) may a sample size bias.

**Fig 10.**
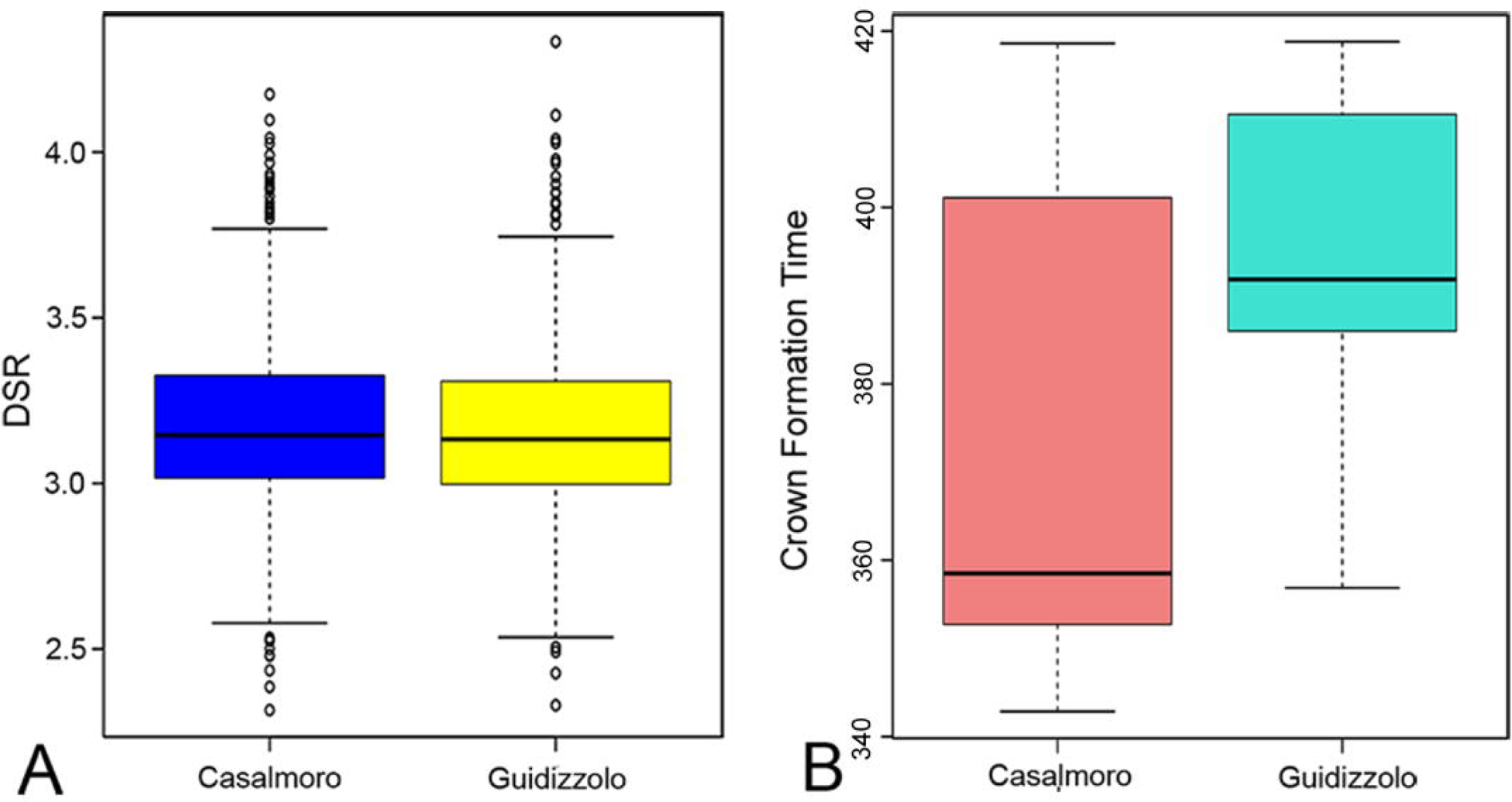
Boxplots of DSR and CFT variation between two sites. Boxplots compare (**A**) daily secretion rates and (**B**) Crown Formation Time from the two necropolises. A marginal yet non-significant distinction was identified only in Crown Formation Time.

Differences were observed between dental arches for Cc but not for Ci. As far as time at Ci is concerned, our results indicate that deciduous first molars started their enamel secretion at a mean 172 days (sd =19) before birth, which correspond to approximately 3.5 months after fertilization (Fig 5). Whereas upper first molars reached Cc around one month before lower molars (Fig 6). Ci estimates for both sites are earlier than what previously reported by Mahoney [51,65], who assessed a mean Ci of 113 days before birth, and Birch and Dean [54] who reported a Ci of 140 days before birth. However, our result fall within the range of variability reported by Lunt and Law [93] and Hillson [94], that is comprised between 3.5 and 4.3 months after fertilization (S5 Table). Cc estimates also differ from Mahoney [51,65] (275 days), falling however within the range of variability reported in Lunt and Law’s review [95] (165-182 days) and being close to the findings of Birch and Dean [54], that found a Cc of 180 days in their sample (S6 Table). The most evident outlier observed in the boxplots (Fig 6) is represented by Guid T.61 LR, which reaches Cc at 310 days after birth. As the CFT of this specimen aligns with those of the other specimens in the sample, this result is likely associated with a delayed Ci (estimated at 88 days pre-birth; Table 1), or/and potentially with a pre-term birth, leading to an extended period of post-birth development. This is also indicated by the more apical position of the NNL compared to the other teeth (Fig 3). In addition, the presence of a high number of ALs may be associated with a change of DSR during stressful periods [50,71,96], this, in turn, can influence the odontochronology.

Upon examining the enamel growth parameters in relation to sexes, no differences were detected. This suggests that the developmental pace in deciduous teeth is not influenced by sex This findings contrasts with some previous studies, which observe a slight advancement in male dental development [97,98]. However, it is worth noting that differences in skeletal development between sexes are commonly known [99]. It is important to consider that in our samples the sex ratio is skewed toward males, comprising approximately 70% of the individuals. This imbalance in the sample composition may have implications for the interpretation of our findings and suggests further investigation into histological aspects of deciduous teeth considering sex.

Finally, the lack of significant differences in all evaluated parameters between the two necropolises (Fig 10) is consistent with observations made by Ballarini [76] and Menotti [77] regarding the similarity – and potential interrelationship – between these two groups, for which certain details of the burial customs set them apart from other allochthonous groups within the same geographic area. This observation warrants further and broader investigations that could provide valuable insight into population dynamics in Northern Italy plains during the Early Medieval period.

## Conclusions

This study contributes to the existing knowledge on developmental patterns of first deciduous molars by collecting data with high temporal accuracy, enabling the assessment of both prenatal and postnatal enamel formation within two archaeological populations from Early Medieval northern Italy. The development of a new regression formula tailored to first deciduous molars from a pre-industrial population enhances our ability to extract valuable information concerning odontogenic trajectories. This proves particularly useful in cases where enamel microstructure might be compromised or not clearly visible [8]. Consequently, it facilitates a more accurate reconstruction of early life developmental histories, and potentially enhances accurate age-at-death estimations for infant individuals.

Furthermore, this work expands the existing body of research on growth rates in posterior deciduous teeth, complementing previous research focused on anterior and permanent dentition [55,56]. Prior studies on first deciduous molars have reported differing developmental patterns both across and within populations and historical periods [35,55,56]. These variations are also evident when comparing our findings with those derived from British Medieval populations [51,54]. Consequently, this emphasizes the need to broaden research to encompass diverse populations from various regions and historical periods. Such an approach is instrumental in acquiring a comprehensive understanding of odontogenic variability, and it aids in establishing geographical and historical standards able to enhance the accuracy of anthropological and archaeological investigations reliant on dental developmental patterns, highlighting the importance of further investigation to attain more accurate and reliable data for deciduous teeth.

In conclusion, employing a multi-analytical approach integrating histological analysis with sex determination through peptide assessment, suggests a potential expansion of histological analysis in archaeological deciduous teeth. This highlights the possibility of conducting furthers sex-driven investigations, particularly concerning the histology of infant in past populations.

## Acknowledgements

We acknowledge the Soprintendenza Archeologica, Belle Arti e Paesaggio per le Province di Cremona, Lodi e Mantova for granting us the permission to conduct the analysis. The authors thank the ‘Fondazione Cassa di Risparmio di Modena’ for funding the nLC-MS system at the Centro Interdipartimentale Grandi Strumenti of UNIMORE. Dr. Diego Pinetti and Dr. Filippo Genovese are thanked for the assistance during LC-MS analyses.

## Supporting Information

**S1 Fig. Chromatograms of amelogenin isoforms in individuals from Casalmoro.** The presence of both isoforms, AMELX (time ∼30 min) and AMELY (time ∼20 min), determines sex as male. Conversely the presence of the only AMELX isoform assesses the sex as female. From Casalmoro 3 individuals are estimated as male and 5 as female.

**S2 Fig. Chromatograms of amelogenin isoforms in individuals from Guidizzolo.** The presence of both isoforms, AMELX (time ∼30 min) and AMELY (time ∼20 min), determines sex as male. Conversely the presence of the only AMELX isoform assesses the sex as female. From Guidizzolo were identified 5 females and 12 males.

**S3 Fig. Linear models to estimate the crown formation time from prism lengths on first deciduous upper and lower molars.** The regression formula for upper molars is Y = 0.315 X and estimated mean DSR was 3.17 μmday^−1^. The regression formula for lower molars is Y = 0.318 X and estimated mean DSR was 3.15 μmday^−1^.

**S4 Fig. Plots of Enamel Extension Rate during lifetime of each individual.** The plots show changes in EER during the lifetime of each individual. Black lines represent the moment of ALs appearance, which are associated with stress events.

**S5 Table. Crown initiation and Crown completion times in first deciduous molars reported in the literature.** Ci times are reported in months after fertilization (considering a gestation length of 39 weeks), whereas Cc times are reported in months after birth.

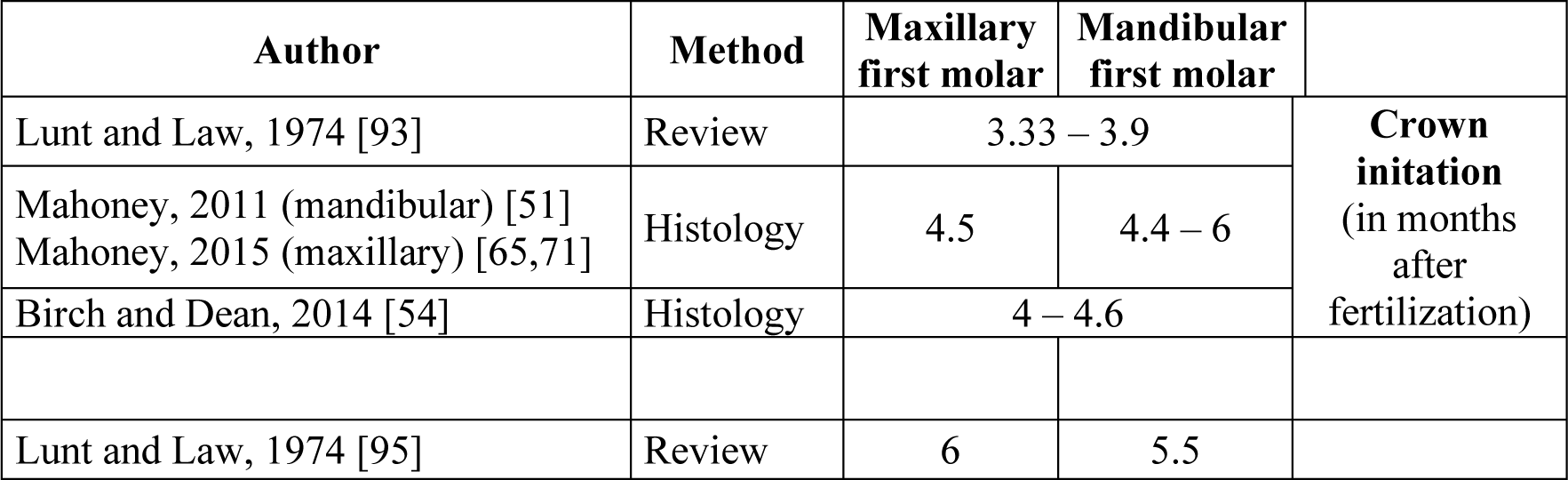
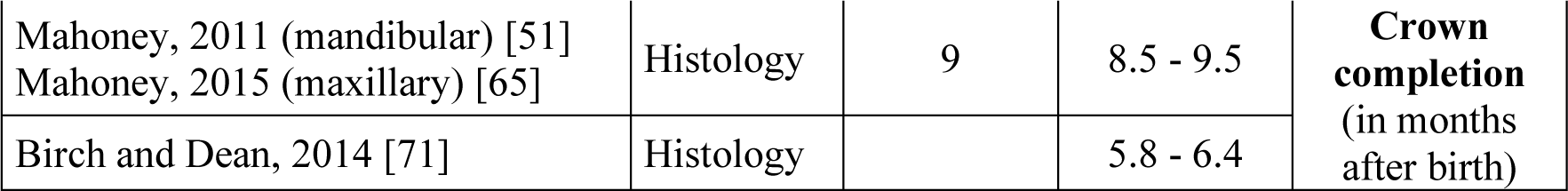

