## Supplementary figures and images for "Enamel histomorphometry, growth patterns and developmental trajectories of the first deciduous molar in an Italian early medieval skeletal series"

### Supplemental Figure 1

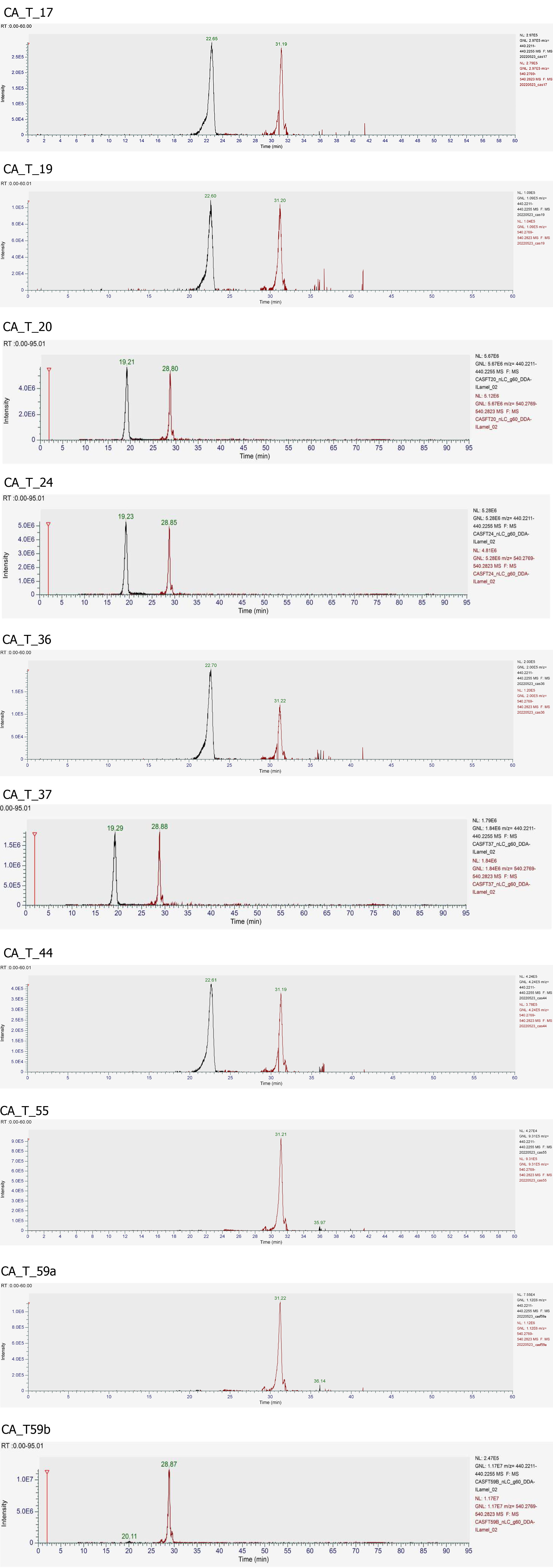

### Supplemental Figure 2

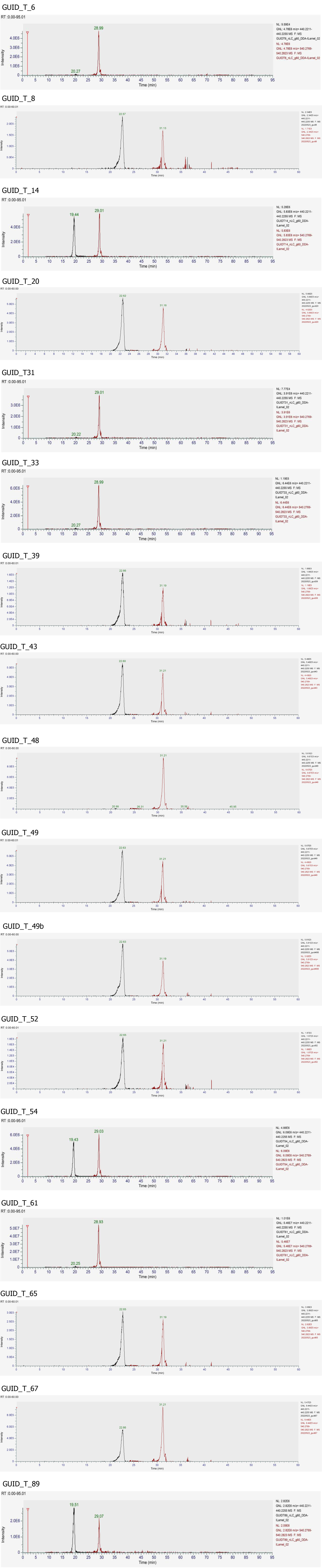

### Supplemental Figure 3

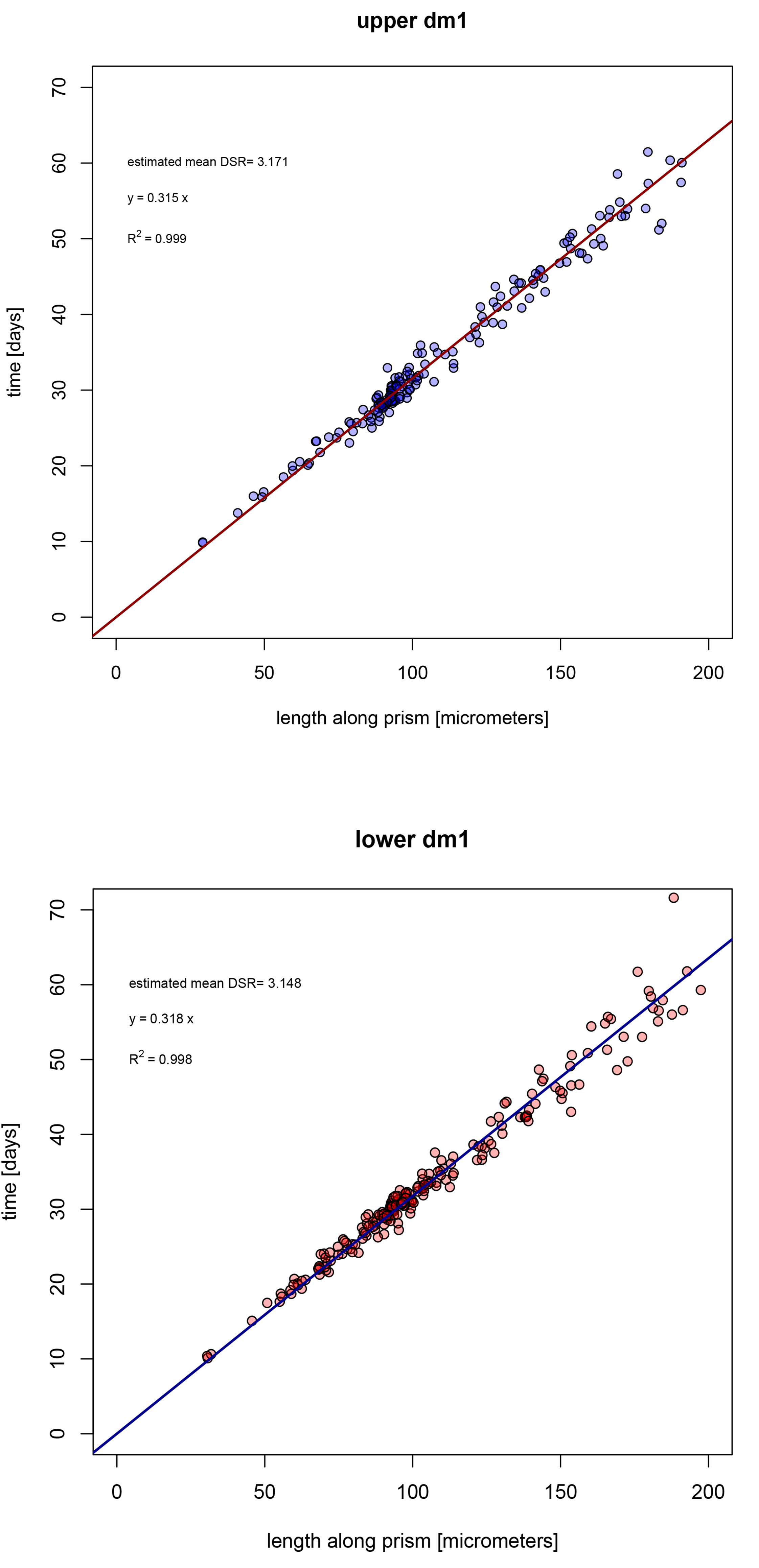

### Supplemental Figure 4

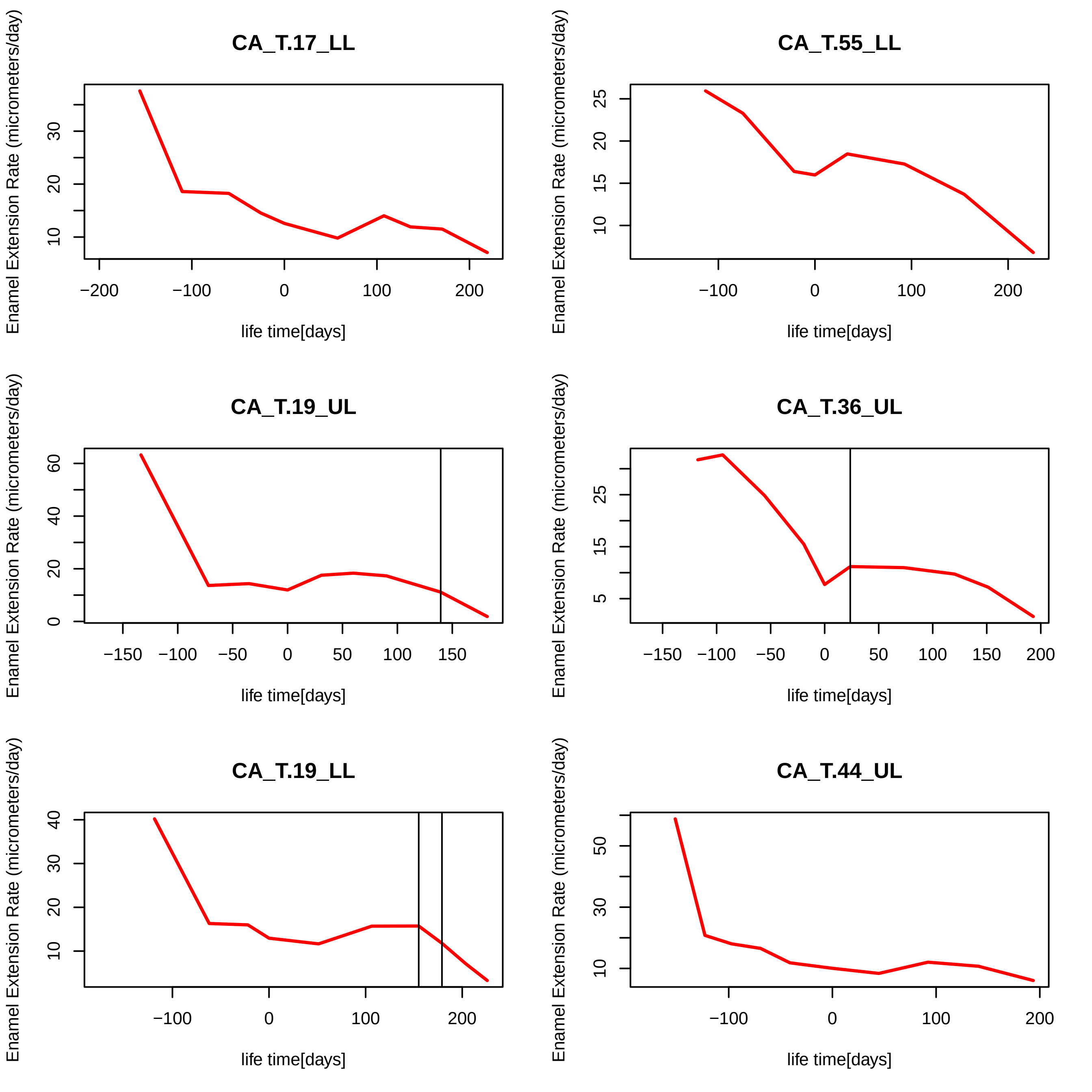

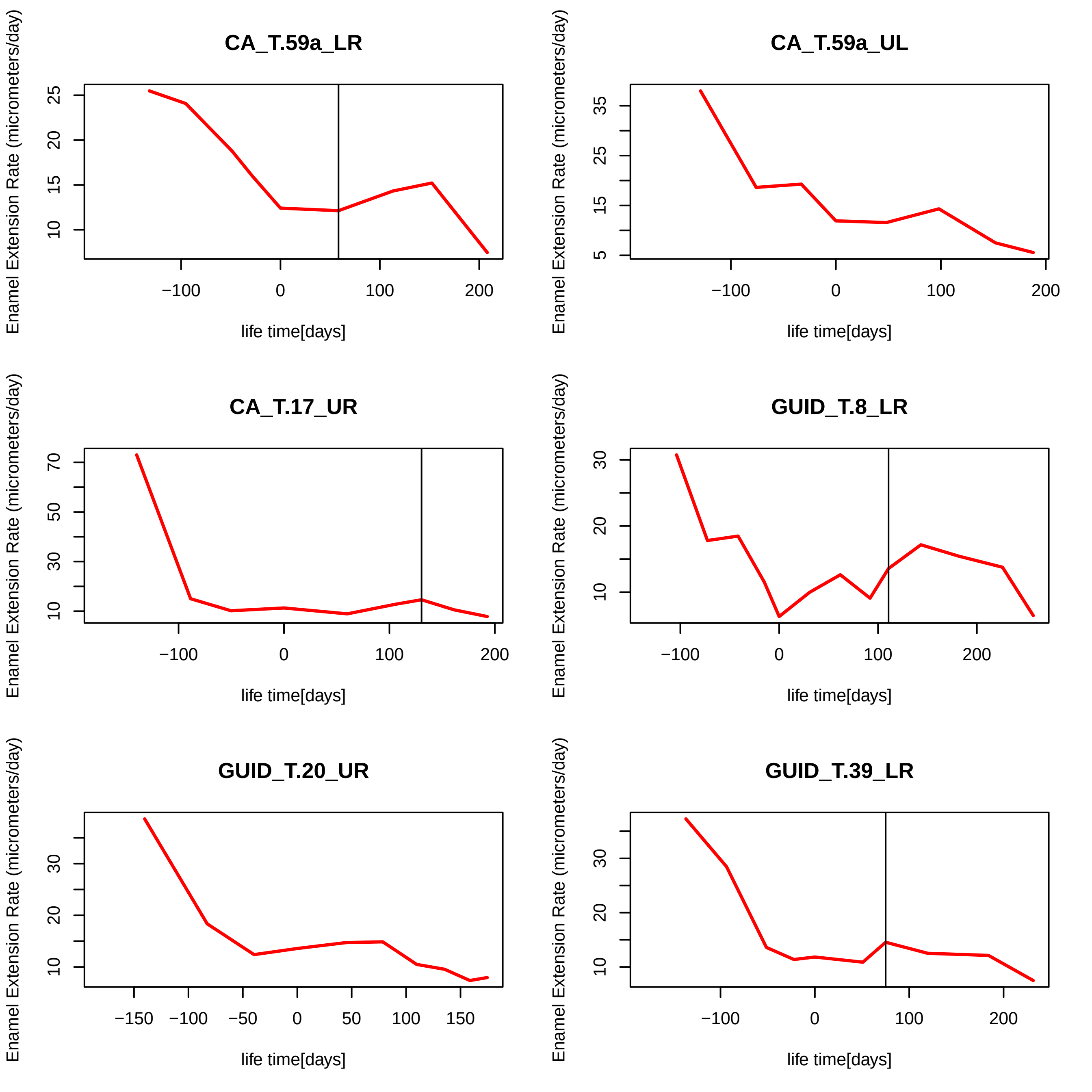

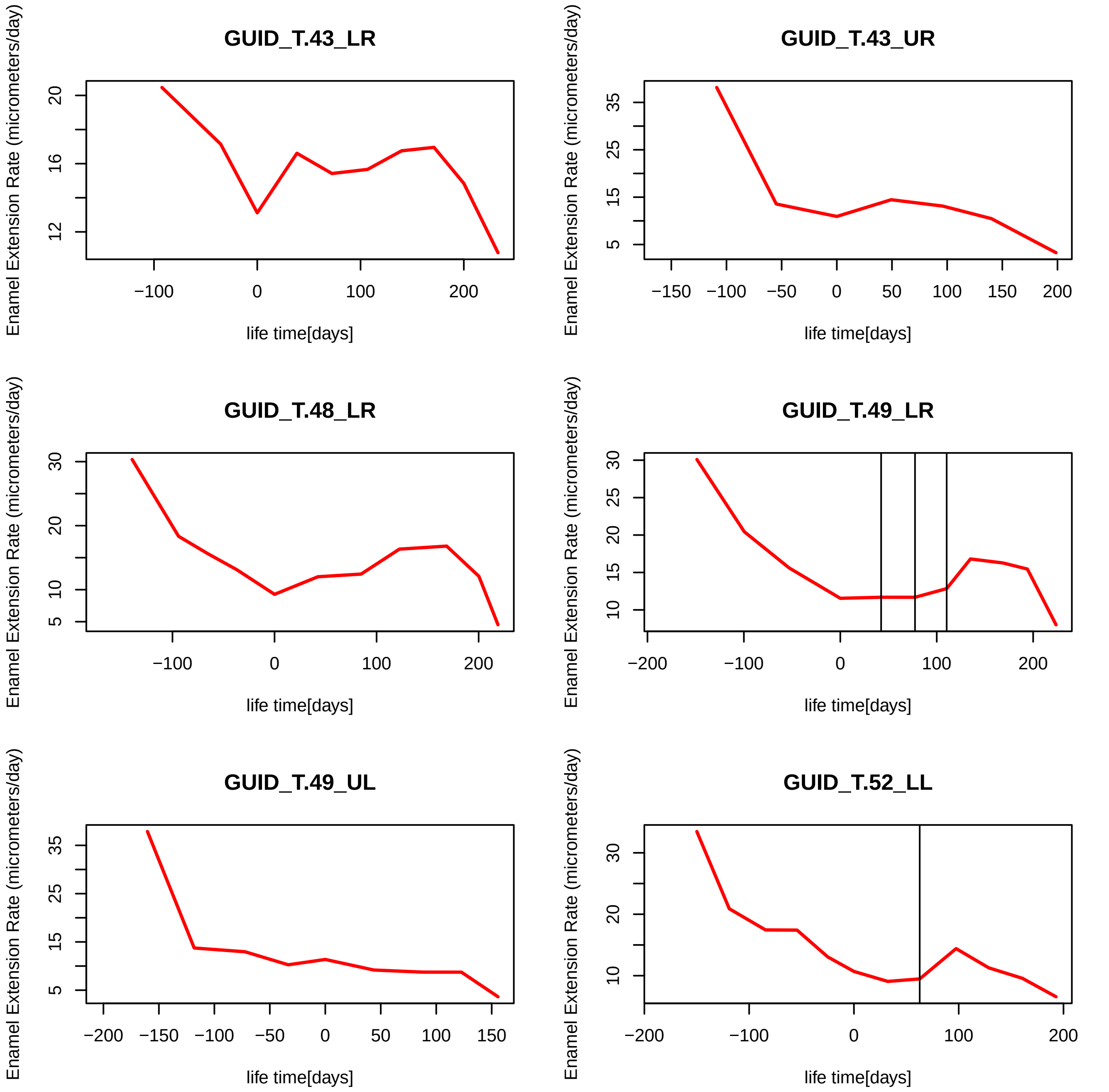

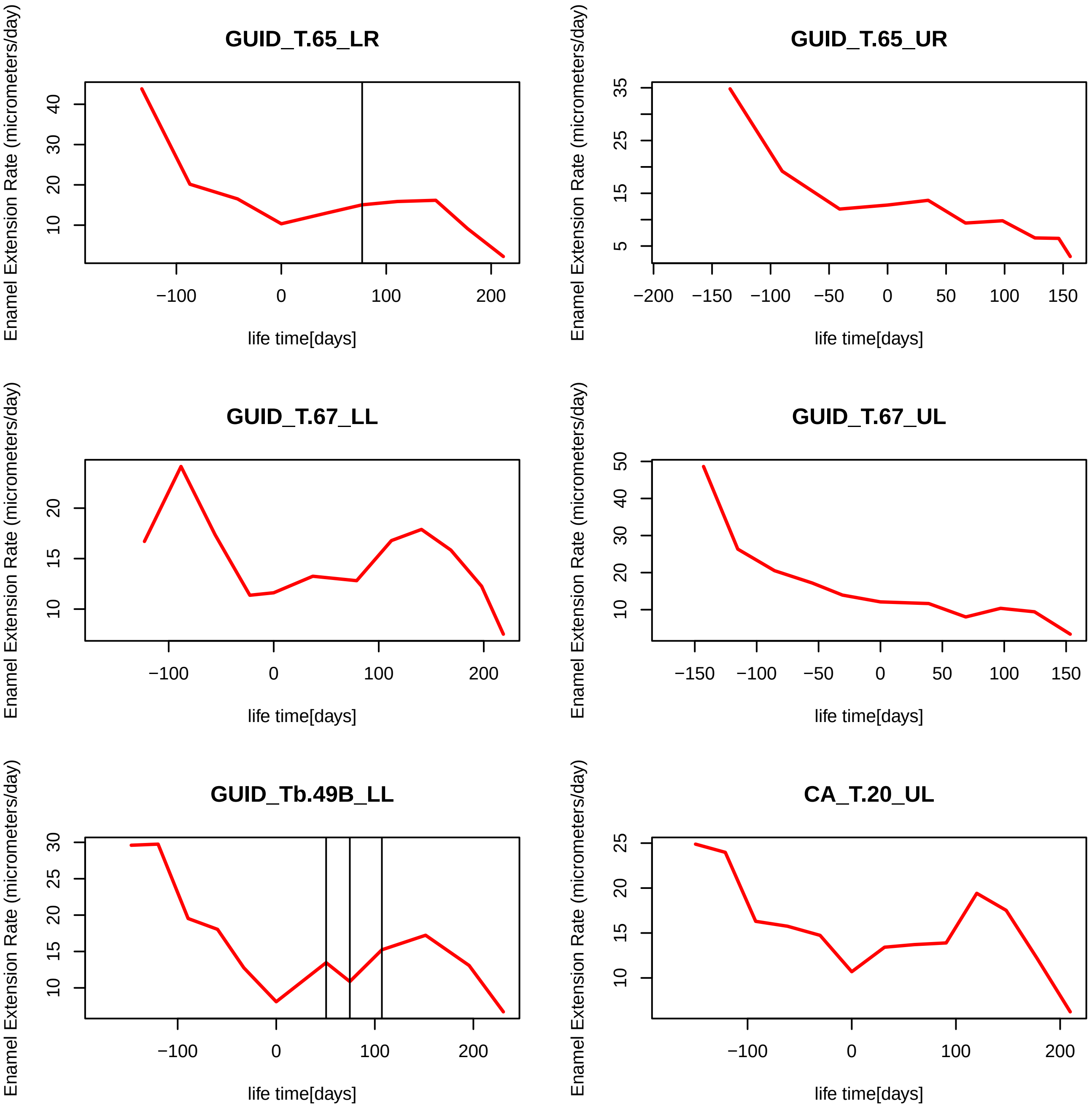

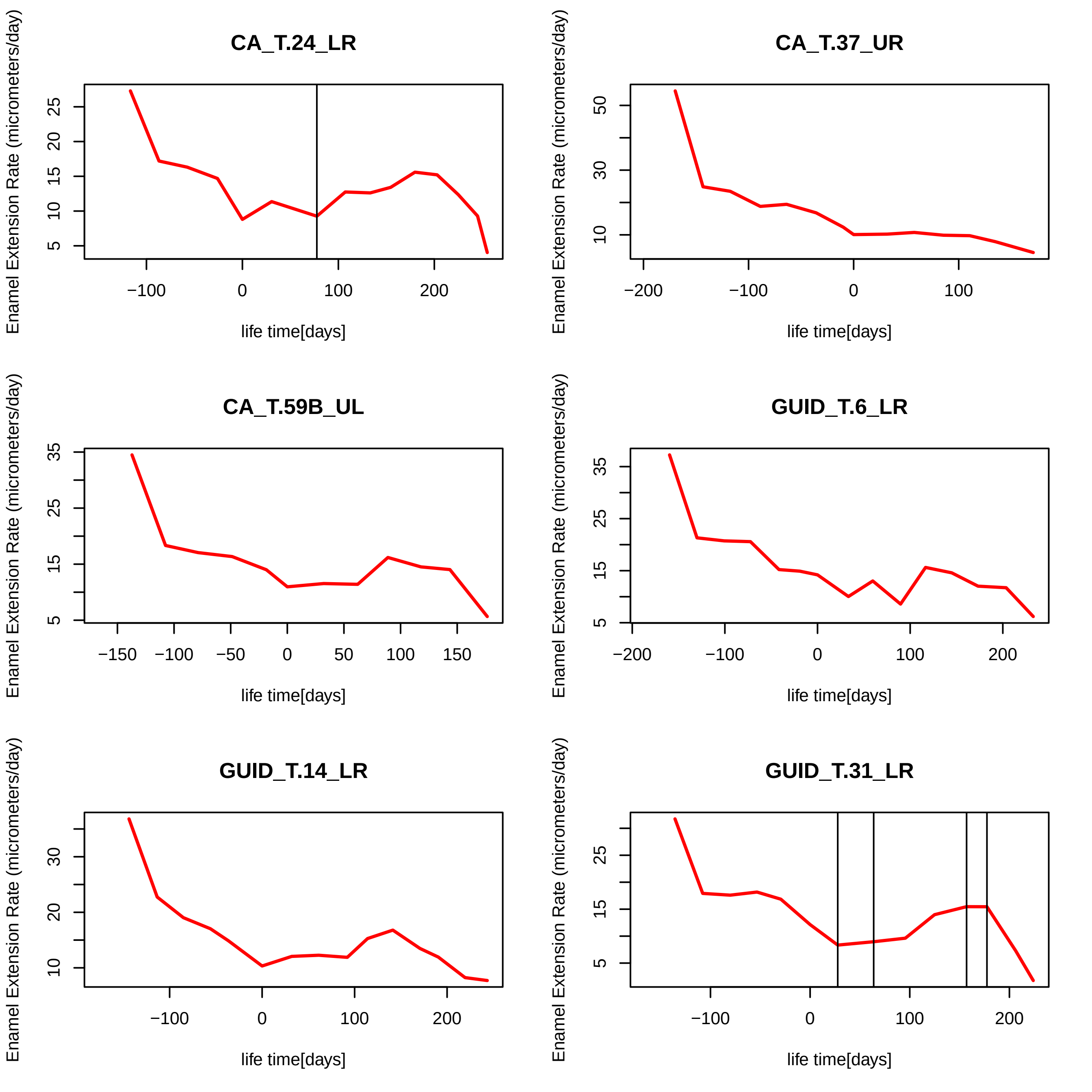

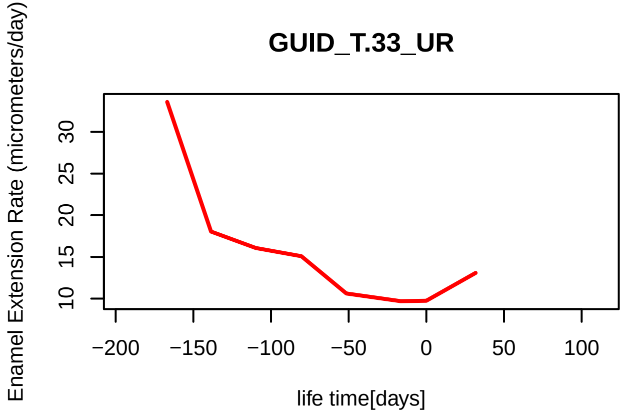

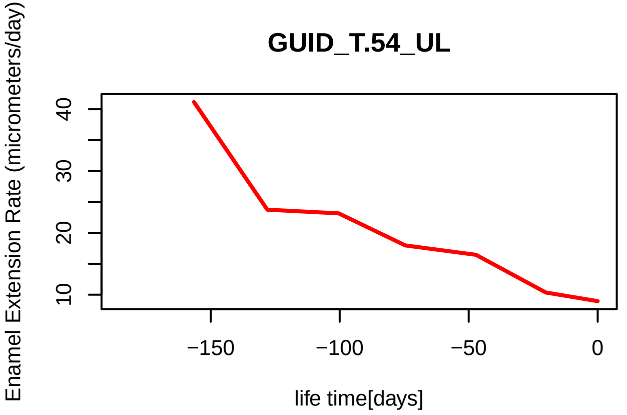

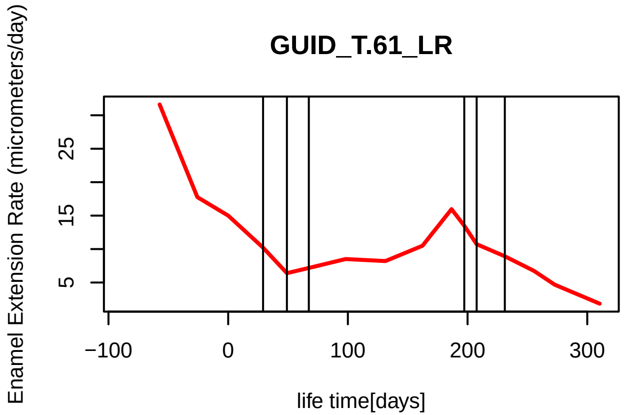

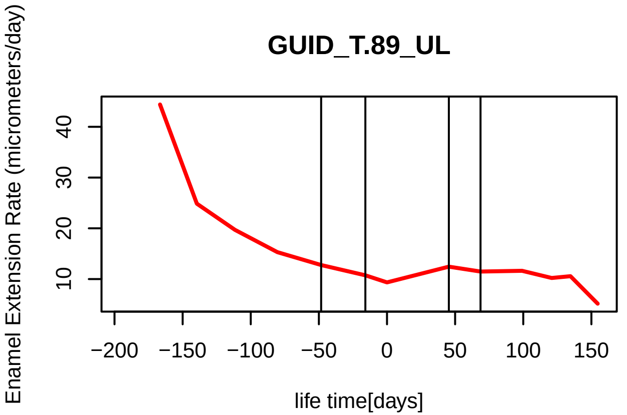
